# Neuronal activity inhibits axonal mitochondrial transport in a region-specific manner

**DOI:** 10.1101/2024.06.04.597338

**Authors:** Venneman Tom, Pieter Vanden Berghe

**Affiliations:** Lab for Enteric NeuroScience (LENS), O&N 1, # 701 Targid, Herestraat 49, KU Leuven, Belgium

## Abstract

Due to their large scale and uniquely branched architecture, neurons critically rely on active transport of mitochondria in order to match energy production and calcium buffering to local demand. Consequently, defective mitochondrial trafficking is implicated in various neurological and neurodegenerative diseases. A key signal regulating mitochondrial transport is intracellular calcium. Elevated Ca^2+^ levels have been demonstrated to inhibit mitochondrial transport in many cell types, including neurons. However, it is currently unclear to what extent calcium-signaling regulates axonal mitochondrial transport during realistic neuronal activity patterns. We created a robust pipeline to quantify with high spatial resolution, absolute Ca^2+^ concentrations. This allows us to monitor Ca^2+^ dynamics with pixel precision in the axon and other neuronal compartments. We found that axonal calcium levels scale with firing frequency in the range of 0.1-1µM, whereas KCl-induced depolarization generated levels almost a magnitude higher. As expected, prolonged KCl-induced depolarization did inhibit axonal mitochondrial transport in primary hippocampal neurons. However, physiologically relevant neuronal activity patterns only inhibited mitochondrial transport in axonal segments which made connections to a target neuron. In ‘non-connecting’ axonal segments, we were unable to trigger this inhibitory mechanism using realistic firing patterns. Thus, we confirm that neuronal activity can indeed regulate axonal mitochondrial transport, and reveal a spatial pattern to this regulation which went previously undetected. Together, these findings indicate a potent, but localized role for activity-related calcium fluctuations in the regulation of axonal mitochondrial transport.

## INTRODUCTION

Though the brain only makes up 2% of our body by weight, it consumes 20% of our total resting energy production^1^. This energy is mainly produced by mitochondria in the form of ATP through oxidative phosphorylation^2^. Remarkably, a single cortical neuron is estimated to consume around 4.7 billion ATP molecules per second^3^. Most of this energy is spent on reversing the ion influxes that underlie synaptic signaling and action potential (AP) firing^4^. Additionally, mitochondria play a key role in calcium homeostasis and signaling, by sequestering Ca^2+^ influx and buffering [Ca^2+^]_i_ fluctuations^2,5^. Due to their large scale and uniquely branched architecture, neurons critically rely on active transport of mitochondria in order to match energy production and calcium buffering to local demand^6^. Consequently, defective mitochondrial trafficking is implicated in various neurological^7–9^ and neurodegenerative diseases^10,11^.

Long-range mitochondrial transport occurs along microtubule tracks. In the axonal compartment, these cytoskeletal polymers are uniformly oriented plus-end out. Therefore, kinesin motors exclusively power anterograde axonal transport, while dynein motors propel cargo in the retrograde direction^12^. Hence, time-lapse imaging of mitochondrial transport in single axons is a powerful method to study transport regulation, enabling direct quantification of effects on either family of motors^13^. These molecular motors are supported by various adaptors, facilitating regulation^14^. The result is a highly complex and dynamic system, characterized by ‘saltatory’ movement; mitochondria frequently pause, change velocity and switch directions^15^.

A key signal regulating mitochondrial transport is intracellular calcium. Elevated Ca^2+^ levels have been demonstrated to inhibit mitochondrial movement in many cell types, including neurons^16–19^. Two molecular mechanisms have been identified so far; one involves an adaptor complex, called the MIRO1-TRAK2-kinesin complex^18–20^, another involves a static anchor protein, syntaphilin^21,22^. According to popular theory, these mechanisms recruit mitochondria to sites with increased Ca^2+^ buffering and energy demands, by inhibiting movement where Ca^2+^ levels are elevated^14^. However, these inhibitory mechanisms were predominantly demonstrated using sustained [Ca^2+^]_i_ elevations, e.g. following a 3-10min exposure to KCl, calcimycin or glutamate^16–22^. Indeed, Ca^2+^ levels can spike a 100-fold during neuronal activity; however, they are rarely sustained under physiological conditions^23^. Hence, it is currently unclear to what extent calcium-signaling regulates axonal mitochondrial transport during realistic neuronal activity patterns.

In this study, we confirm that neuronal activity can indeed regulate axonal mitochondrial transport, but we reveal a spatial pattern to this regulation which went previously undetected. Prolonged KCl-induced depolarization of the neuron did inhibit axonal mitochondrial transport in line with previous reports. However, realistic neuronal activity patterns only inhibited mitochondrial transport in axonal segments connected to a target neuron, not in segments without these connections. Together, the findings presented in this paper indicate a potent, but localized role for activity-related calcium signaling in the regulation of axonal mitochondrial transport.

## RESULTS

### Activity-related calcium spikes are not sufficient to halt axonal mitochondrial transport

To investigate to what extent Ca^2+^ fluctuations impact axonal mitochondrial transport under physiologically relevant conditions, we optimized a set-up to simultaneously record mitochondrial transport and calcium dynamics in single axons of primary hippocampal neurons in sparse cultures (fig. 1a). Axon origin and identity were verified via post hoc immunolabeling (fig. 1a, inset), enabling the distinction between anterograde and retrograde transport. To induce the firing patterns of choice, precise electrical field stimulation (EFS) was performed by carefully positioning a custom-made bipolar electrode around the cell soma (fig. 1b). Time-lapse imaging at high temporal resolution (∼300Hz), using the voltage sensor BeRST1^24^ in combination with the calcium indicator Fluo-4, confirmed the tight coupling between action potential firing and calcium spikes (fig. 1c-d).

**Figure 1.**
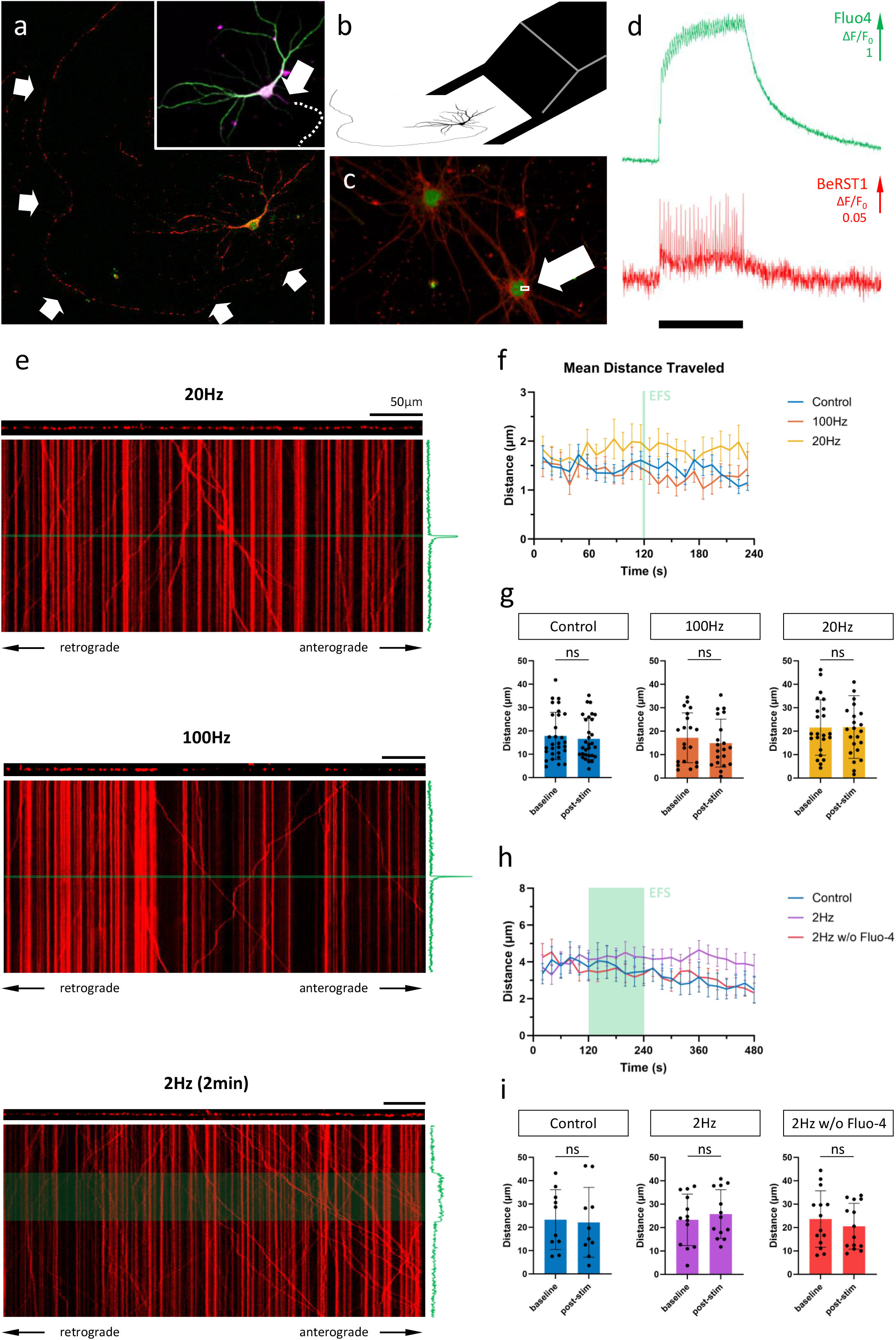
Axonal mitochondrial transport is unaffected by neuronal activity patterns. (a) Representative field of view (FOV) for near-simultaneous recording of calcium activity (Fluo-4) and axonal mitochondrial transport (mitoTracker Red). Arrows mark axonal fiber. Inset: MAP2 (green), AnkG (magenta) and Tau immunolabeling to verify axonal identity and origin. Arrow indicates axon initial segment. Dotted line indicates axonal path. (b) Schematic representation of bipolar electrode positioning. (c) Primary hippocampal neurons loaded with cytosolic calcium indicator Fluo-4 (green) and membrane-bound voltage sensor BeRST1 (red). Arrow indicates 4×2 pixel ROI used for high-frequency scanning. (d) Calcium and voltage responses during 20Hz stimulation (black bar marks stimulation period = 2s). (e) Kymographs of representative recordings of the 3 main stimulation paradigms; 20Hz 2s, 100Hz 2s and 2Hz 2min. The axonal segment is shown above each kymograph and the axonal Fluo-4 signal along the time axis. Green line/box indicates stimulation period. Scalebars = 50µm. (f) Binned distance over time reveals no effect on transport during/following activity. (g) Quantification of distance traveled during baseline period (1^st^ half of recording) vs. post-stimulation (2^nd^ half of recording). Mann-Whitney U test, p = 0.62 (ctr), 0.49 (100Hz), 0.96 (20Hz), n = 30 rec & 255 mob mito (ctr), 20 rec & 145 mob mito (100Hz), 23 rec & 205 mob mito (20Hz). (h-i) idem panel f-g. For these recordings, the first 2min (= baseline) were compared to the 2min stimulation period, with and without fluo-4 present. Mann-Whitney U test, p = 0.74 (ctr), 0.48 (2Hz), 0.60 (2Hz w/o Fluo-4), n = 10 recs, 141 mob mito (ctr), 13 recs, 196 mob mito (2Hz), 14 recs, 189 mob mito (2Hz w/o Fluo-4).

Since KCl-induced depolarization has been used previously to demonstrate the calcium-dependent inhibition of mitochondrial transport, we first repeated this finding (fig. 2a-c). In line with previous reports, a 2-minute KCl perfusion resulted in a substantial, yet temporary, decrease in axonal mitochondrial transport (fig. 2d-e). Such a sustained depolarization, however, does not perfectly represent the calcium spikes which occur during neuronal activity. Therefore, to more closely match physiological conditions, we then tested how activity patterns of different frequencies affected transport. Analysis of transport parameters, such as distance traveled (fig. 1f-i), mobile fraction, motor velocity, start & stop probabilities and run length, was performed based on manual tracings of mitochondrial trajectories in kymographs, 2D projections of the axonal segment (fig. 1e). Surprisingly, none of these movement parameters were affected by the activity patterns that were tested (fig. 1f-g), even at the highest stimulation frequency (100Hz). To assess whether the inhibitory effect scales with the duration of the activity pattern, rather than the frequency, we tested another stimulation paradigm during which cells were induced to fire for an extended period of time (2min, 2Hz). Again, there was no effect on the distance traveled by mitochondria (fig. 1h-i), or any of the other transport parameters (not shown). To rule out the potential chelation of calcium by the indicator itself, the experiment was repeated in the absence of a calcium indicator. Likewise, axonal mitochondrial transport, in either direction, remained unaffected by the induced activity patterns (fig. 1 h-i).

**Figure 2.**
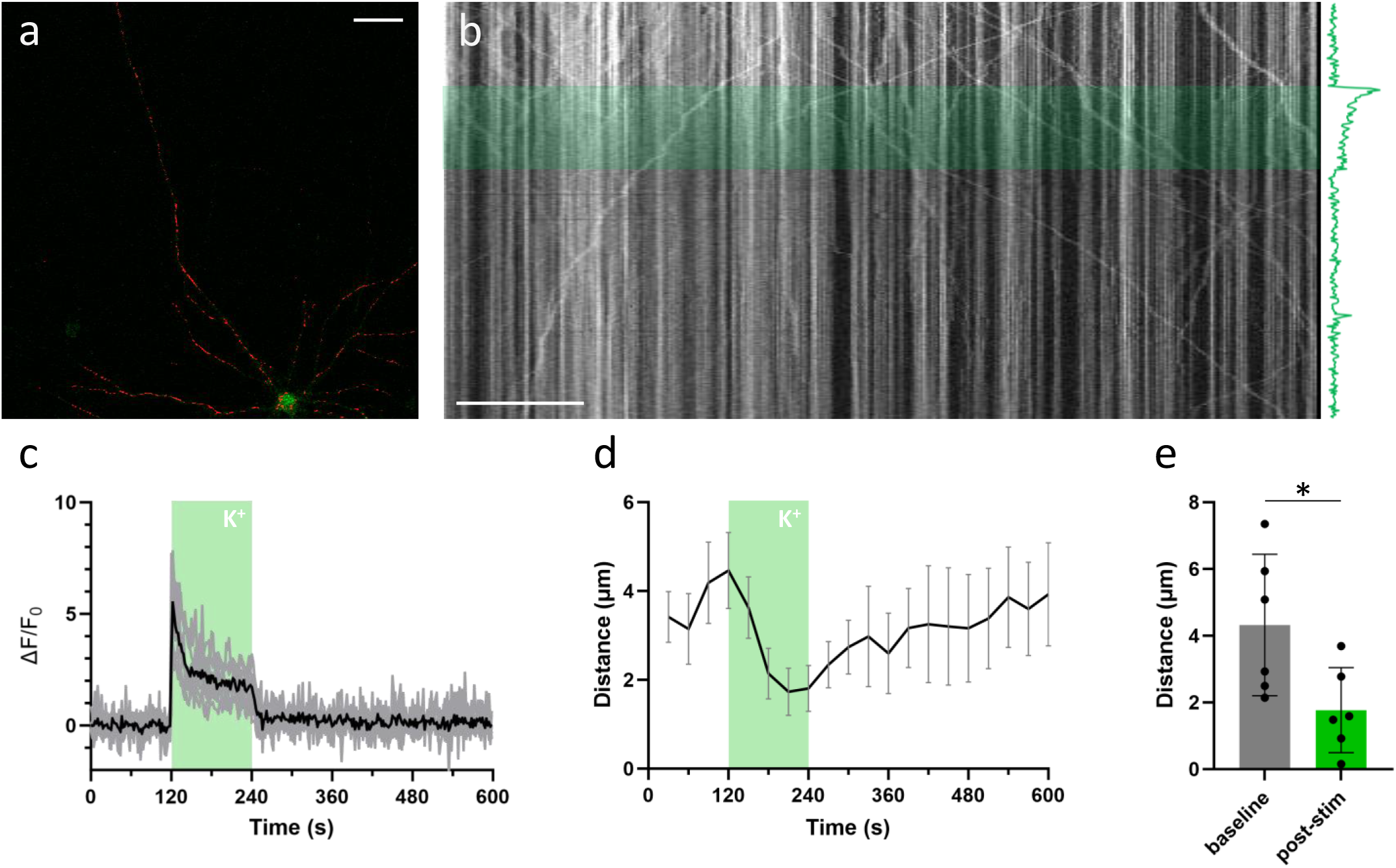
KCl perfusion inhibits axonal mitochondrial transport. (a) Example neuron, loaded with mitoTR CMXRos and Fluo4 (Scalebar = 50µm). (b) Kymograph from axonal segment. The KCl perfusion period (2min) is marked in green. The axonal Ca^2+^ response is shown along the time-axis. (c) Mean axonal Ca^2+^ response to KCL perfusion in black, individual traces in gray. (d) Axonal transport over time (quantified as the mean distance traveled per mitochondrion, binned every 30s. Errorbars = SEM). KCl perfusion period (2min) is marked in green. (n = 6 recs, 8 segm, 97 mob mito). (e) Statistical comparison of mean distance traveled between baseline period (minute before stimulation) vs. last minute of stimulation period. Mann-Whitney U test (p = 0.04). Errorbars = STD.

### Axonal calcium levels scale with firing frequency in the range of 0.1-1µM

In contrast to KCl-induced depolarization (fig. 2), activity-induced calcium elevations did not inhibit axonal mitochondrial transport (fig. 1). A potential explanation might be that the calcium spikes which occur during spontaneous firing (and EFS-induced activity patterns) are insufficiently high to trigger this inhibitory effect in the axon. However, to our knowledge, no method is available to measure the axonal Ca^2+^ dynamics with pixel precision. To this end, we created a robust method to quantify absolute Ca^2+^ concentrations in neuronal compartments with high spatial precision, using the ratiometric Ca^2+^ indicator Fura-2 (fig. 3a).

**Figure 3.**
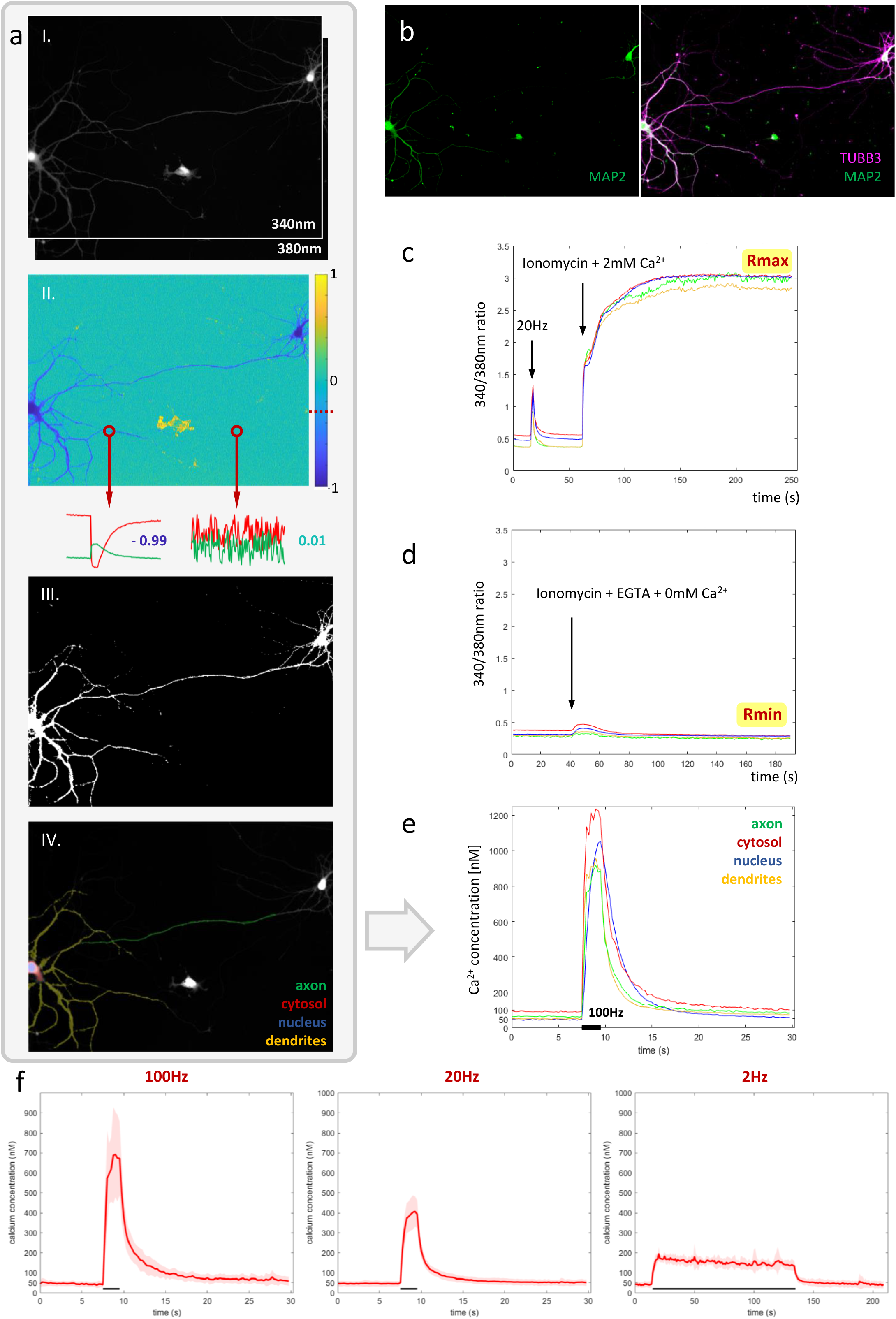
A robust pipeline for measuring absolute calcium concentrations in the axon and other neuronal compartments. (a) Key steps in the pipeline. From top to bottom: (I.) Near-simultaneous widefield imaging at 340 & 380nm, (II.) inverse correlation-based thresholding, (III.) resulting neuronal mask, (IV.) used-defined compartment-ROIs. (b) Immunolabeling for MAP2, TUBB3 and DAPI to distinguish compartments: axon, dendrites, soma and nucleus. (c-d) Calibration of 340/380nm ratio to absolute calcium concentration in living neurons, using ionomycin to saturate and deplete intracellular calcium. (e) Absolute calcium responses measured in each neuronal compartment for representative recording of a primary hippocampal neuron shown in panel a, stimulated at 100Hz for 2s. (f) average axonal responses to 3 main stimulation paradigms. n = 6 cells (100Hz), 13 cells (20Hz), 6 cells (2Hz).

First, image stacks are registered and background-corrected. Next, we exploit the tight inverse correlation between the 340 & 380nm signals exhibited by pixels within the responding cell, to segment a mask of the neuron (fig. 3a). Compared to intensity-based thresholding, this method enables near-perfect separation from non-responsive cells or fibers, moving cells and debris (fig. 5). The resulting binary mask is used to define compartment-specific ROIs, guided by post-hoc immunolabeling using anti-MAP2 and anti-TUBB3 antibodies (fig. 3b). Since ratiometric methods are sensitive to background contributions, we also corrected the extracted ROI-signals for their compartment-specific autofluorescence contribution, which was measured in unloaded neurons (not shown).

Finally, absolute calcium concentrations were calibrated based on a set of reference measurements, performed in primary hippocampal neurons under identical conditions. Briefly, the maximal and minimal 340/380nm ratios (R_max &_ R_min_) were determined by intracellular measurement in living neurons, using ionomycin and varying concentrations of external Ca^2+^ to saturate and deplete intracellular calcium levels, respectively (fig. 3c-d).

To account for differences in excitation (340 & 380nm) and efficiency of the Ca^2+^-bound and Ca^2+^-free Fura2-signals, a ROI or compartment-specific correction value alpha (α) is used to calculate the effective dissociation constant K_eff_:

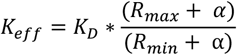

Which is used to calculate the actual intracellular Ca^2+^ concentration, using:

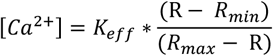

This approach enabled us to monitor Ca^2+^ dynamics in great detail, within different compartments of the same neuron (fig. 3e). A ∼50nM baseline was observed in all compartments, except for the cytosolic one, where values ranged between 50-100nM (fig. 4). Calcium responses differed as well between compartments. Ca^2+^ responses typically rose more slowly in the nucleus than in the cytosol, where the concentration reached higher levels than anywhere else in the neuron. A high degree of variability in response amplitude was observed within the dendritic tree, compared to low variability in the axonal response. Figure 3f shows the average absolute calcium responses in the axonal compartment to the 3 types of activity patterns used in previous transport experiments: During a 100Hz 2s burst, Ca^2+^ reached 713 ± 211nM, for 20Hz 2s burst, 419 ± 76nM and for a 2Hz 2min period, 182 ± 17nM.

**Figure 4.**
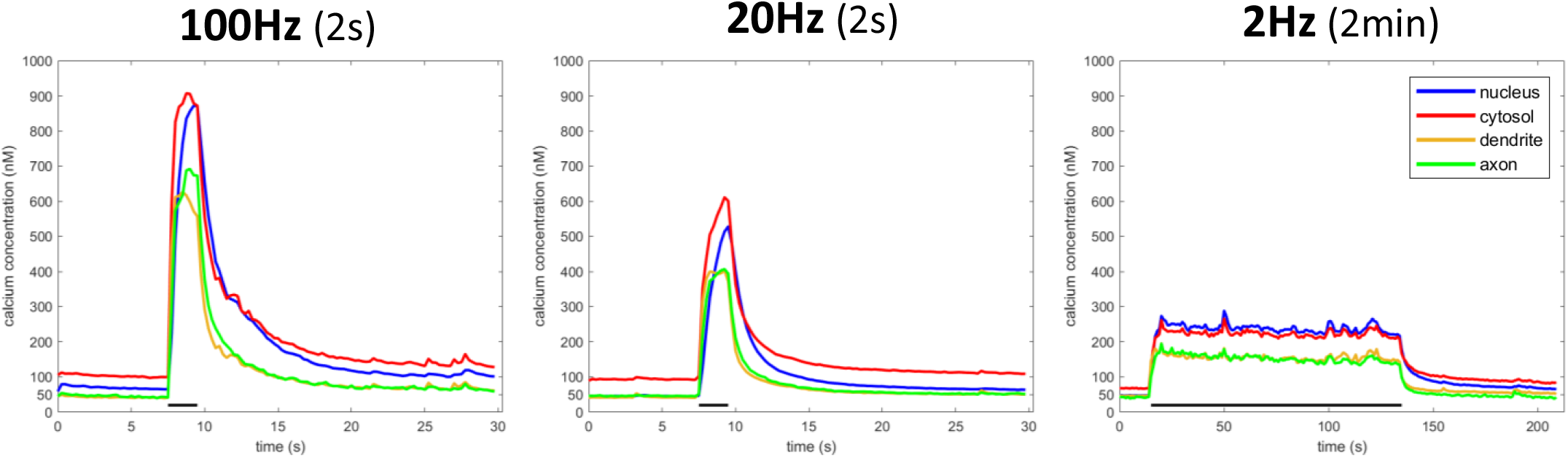
Compartment-specific Ca^2+^ responses to 3 stimulation types. (n = 6 cells (100Hz), 13 cells (20Hz), 6 cells (2Hz))

**Figure 5.**
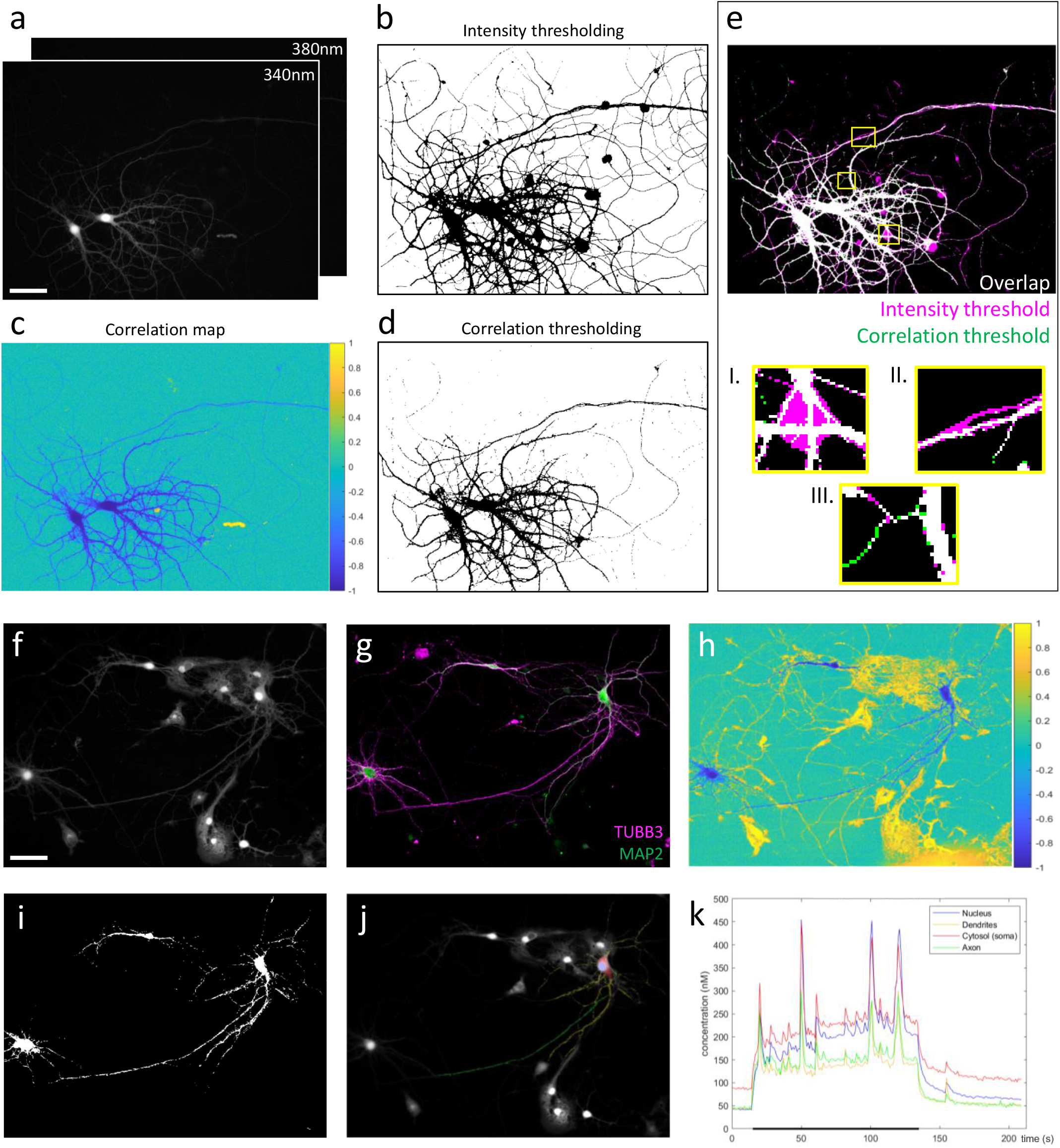
Correlation-based thresholding vs. intensity-based approach for the segmentation of ratiometric image stacks. (a) Raw image stacks (scalebar = 50µm). (b) Segmentation mask obtained by conventional manual thresholding, performed on a median projection of each image stack. The resulting masks were then combined to generate the final mask. (c) Correlation map, generated by pixel-wise correlation of 340 & 340nm pixel traces. (d) Segmentation mask based on correlation thresholding. For all recordings included in the dataset for figure 3, a correlation threshold of -0.3 reliably segmented the neuronal silhouette. (e) Comparison of intensity (magenta) vs. correlation-based (green) segmentation methods (white = overlap). Correlation-based thresholding easily excludes pixels where no reliable measurement can be made and which would otherwise contaminate quantification, e.g. fluorescent debris (I.) and non-responsive fibers (II.). Moreover, responsive yet low-intensity fibers are more easily segmented by correlation-based thresholding owing to their negative correlation values (III.). (f) Example recording that represents near-impossible task for intensity-based segmentation of a neuron surrounded by various other cells (scalebar = 50µm). (g) Immunocytochemistry to distinguish neuronal compartments. (h) Correlation map. Neuron of interest can be identified based on its negative correlation values. Surrounding cells (and debris) earn (neutral or) positive correlation between the 340 & 380nm pixel traces, due to cell movement and signal bleaching. (i) Segmentation mask (threshold = -0.3). (j) User-defined compartmental ROIs. (k) Compartment-specific Ca^2+^ responses during a 2-minute train of action potentials at 2Hz (black bar).

### Activity regulates mitochondrial transport in axonal segments that connect to target neurons

Previous reports have estimated the IC50 value of calcium’s inhibitory effect on mitochondrial transport to be in the range of 400nM^17,18,20^. The firing patterns used in our experiments induced axonal calcium levels exceeding this threshold (fig. 3). Hence, if activity-dependent inhibition of transport was solely dependent on axonal calcium levels, we should have observed a cessation (or substantial decrease) in mitochondrial transport. However, in contrast to KCl-induced depolarization, realistic firing patterns did not inhibit axonal mitochondrial transport (fig. 1), despite inducing sufficiently high Ca^2+^ levels.

Since one of the proposed functions of this regulatory mechanism is to recruit mitochondria to pre^5,22,25^ -and post^20,26^-synaptic locations, we hypothesized that its action might be locally constrained within the axon. Therefore, we set out to test whether neuronal activity affects mitochondrial transport in more distally located segments, where it branches and makes connections to another neuron. To this end, we adjusted our experimental approach. Instead of quantifying transport in extremely sparse cultures in non-connecting (fig. 6a) axonal segments (which lack functional synapses), transport recordings were performed in mixed cultures, made from wildtype and mito-dendra2-positive pups (fig. 6b). This allowed us to quantify mitochondrial transport specifically in axonal segments that connect to a target cell. Post hoc immunolabeling was used to confirm the physical interaction between pre- and post-synaptic cells (fig. 6c-d).

**Figure 6.**
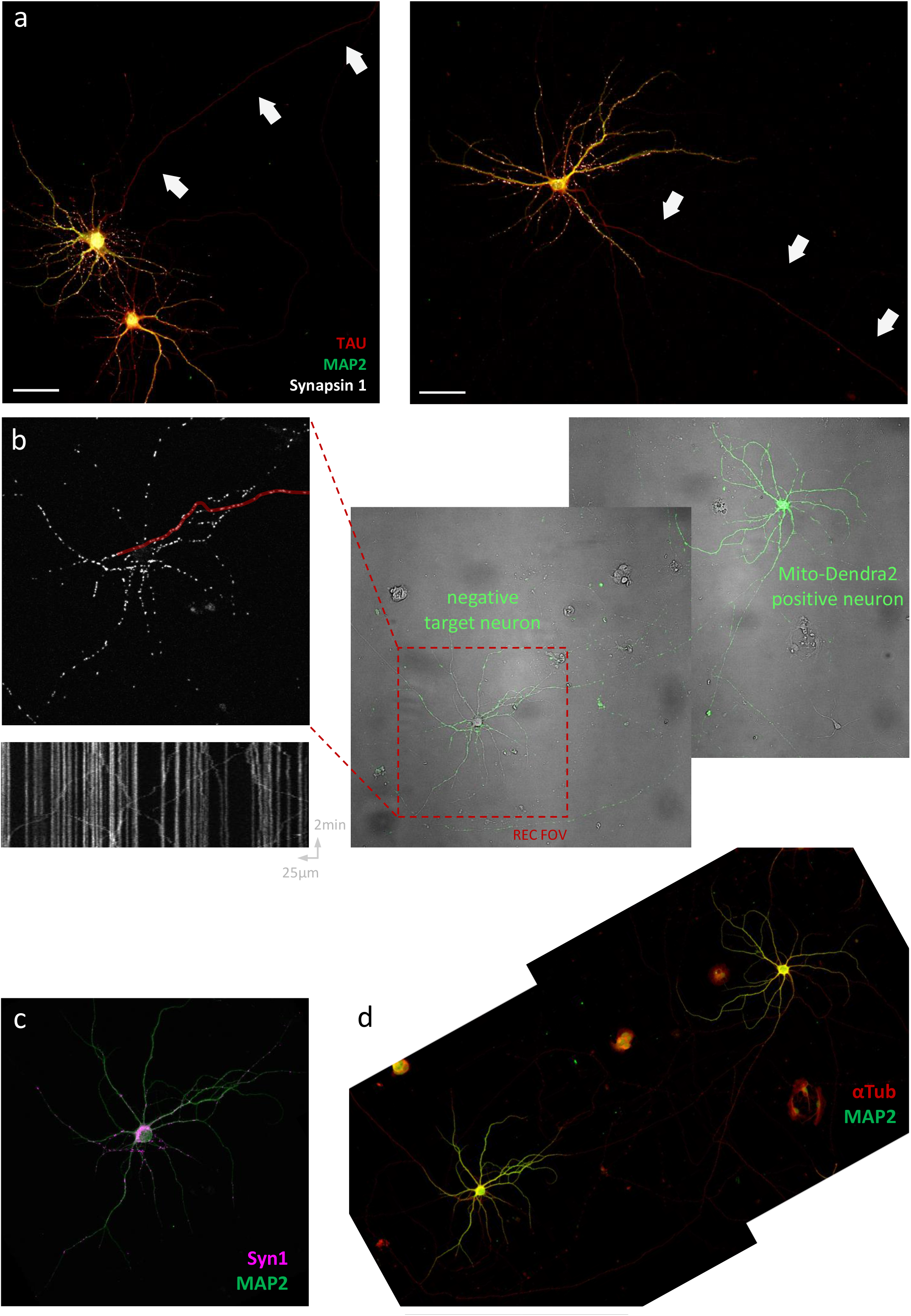
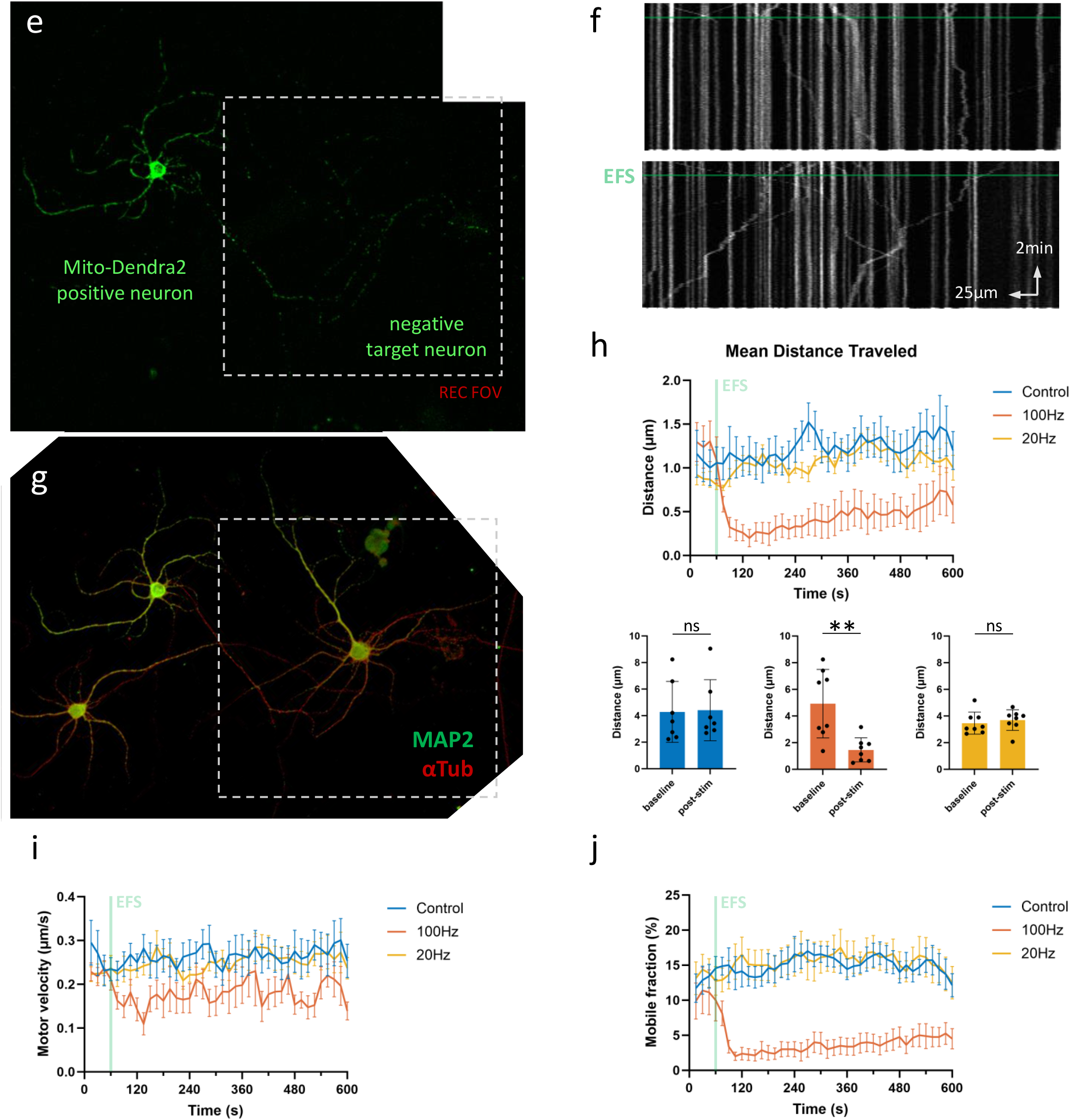
Activity regulates mitochondrial transport in axonal segments that connect to target neurons. (a) Representative FOVs in sparse neuronal cultures. Arrows indicate lack of pre-synaptic sites in non-connecting axonal segments traditionally used for detailed transport measurements. Scalebars = 50µm. (b) Overview protocol to measure axonal mitochondrial transport in connecting axonal segments (control recording, no stimulation). Right to left: mixed cultures of wildtype and mito-dendra2-positive neurons. FOV is chosen at target neuron (red square). Axonal segments are traced (example marked in red) to produce kymographs for analysis. (c-d) Immunolabeling for MAP2 (green), αTub (red) and Syn-1 (magenta) is used to confirm the connection. (e-g) Idem for a representative neuron stimulated at 100Hz for 2s. (f) Kymograph shows mitochondria halting shortly after electrical field stimulation (EFS). (h) Quantification of mean distance traveled over time and statistical comparison between baseline period vs. 1 minute after stimulation. Mann-Whitney U test, p = 0.80 (ctr), 0.0047 (100Hz) and 0.28 (20Hz), n = 7 recs, 21 segm, 360 mob mito (ctrl), 8 recs, 23 segm, 212 mob mito (100Hz), 8 recs, 30 segm, 469 mob mito (20Hz). (i-j) Quantification of velocity & mobile percentage over time.

In these connecting axonal segments, high frequency activity patterns (100Hz, 2s) resulted in a clear inhibition of mitochondrial transport (fig. 6e-h). This decrease was mainly achieved through halting mitochondria rather than slowing down their movements, as evidenced by the peak in pause events and decrease in mobile fraction (fig. 6j), compared to the smaller effect on motor velocity (fig. 6i). All inhibitory effects associated with the stimulation were temporary in nature, and were almost fully recovered by the end of the 10min recording. No effect on transport was observed following a 20Hz (2s) burst of activity (fig. 6h).

## DISCUSSION

Calcium signaling serves a potent role in the regulation of mitochondrial transport and distribution^14^. Since intracellular calcium levels fluctuate during neuronal activity, we set out to study how action potential firing influences axonal mitochondrial transport. Surprisingly, we found no evidence that mitochondrial transport is affected by activity-related calcium fluctuations in non-connecting axonal segments (fig. 1). In axonal segments that do connect to target neurons however, neuronal activity was able to inhibit mitochondrial transport (fig. 6). This spatial pattern might suggest that *in vivo*, neuronal activity similarly does not affect mitochondrial transport along the majority of the projecting axon’s length, but does so only locally, where the axon makes its connection to target cells.

There are likely multiple reasons as to why this pattern was not detected in previous studies. The first is related to culture density. EFS has been used before to demonstrate activity-dependent inhibition of axonal mitochondrial transport^25^. However, we elected to perform our first set of experiments in single axons in highly sparse cultures, where axons can extend for millimeters before connecting to a target neuron. This approach offers key advantages in terms of accessibility for imaging and sample manipulation, as well as analysis strategies. As a consequence, transport recordings biased towards non-connecting axonal segments. As a result, activity-related calcium elevations initially appeared insufficient to halt axonal mitochondrial transport, and finally led us to test a region-specific hypothesis.

Secondly, calcium’s inhibitory effect on mitochondrial transport in neurons has been demonstrated using stimuli of very different proportions. Most often, calcium-dependent inhibition of mitochondrial transport is demonstrated using stimuli, that produce elevations in calcium which surpass those observed during neuronal activity, in both concentration and duration^16–20,22^. Similarly, when we applied KCl-depolarization, we observed a decrease in axonal mitochondrial transport, independent of the presence of a nearby target neuron (fig. 2). However, the axonal calcium levels induced by KCl-depolarization are almost a magnitude higher than those observed during realistic firing patterns (fig. 7).

**Figure 7.**
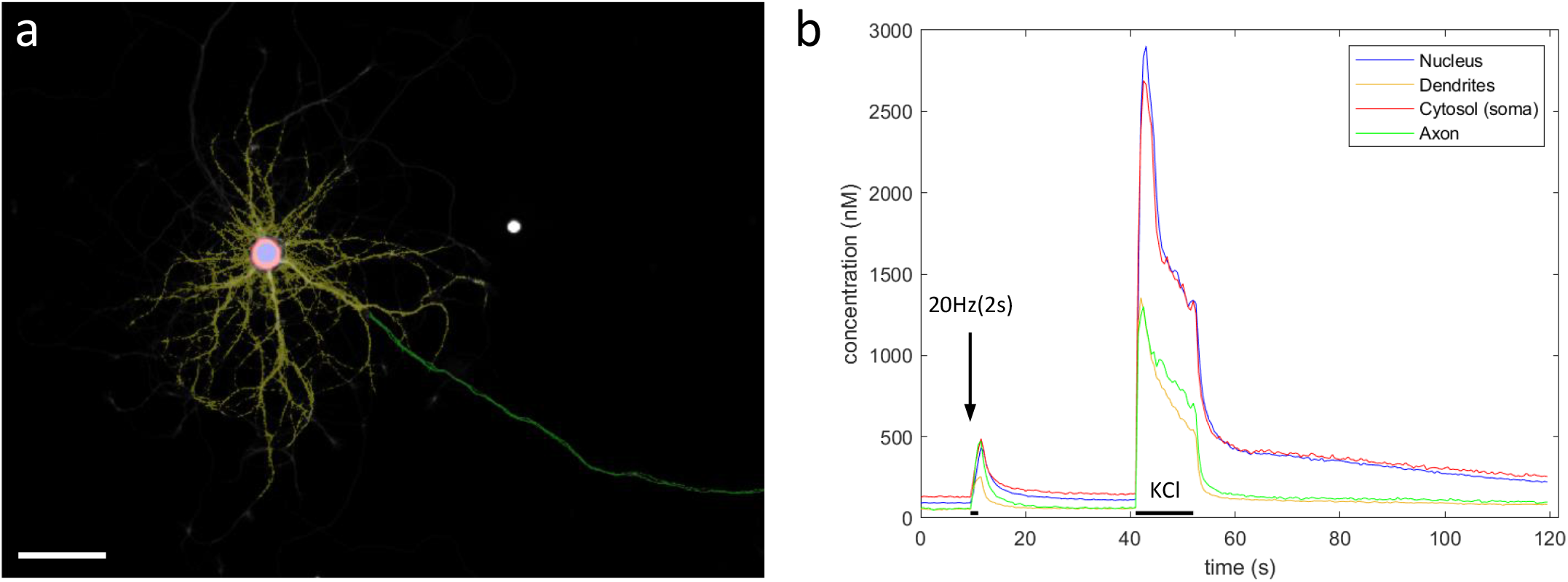
Compartment-specific Ca^2+^ responses to KCl perfusion. (a) User-defined compartmental ROIs (scalebar = 50µm). (b) Compartment-specific Ca^2+^ responses after electrical field stimulation (20Hz 2s) and KCl perfusion (11.5s).

This might suggest the existence of an alternative pathway, which also results in the inhibition of transport, but is triggered by a much higher calcium load. It is possible that such a mechanism could serve a neuroprotective role when calcium levels exceed their physiological range^27^. Indeed, calcium-sensitive adaptors to the motor-complex have been demonstrated to protect against excitotoxicity associated with excessive calcium influx^19^. Alternatively, the cessation of movement under such pathological conditions might result from an indirect effect on mitochondrial function, blocking ATP production. Similarly, calcium overload during epileptic seizures can lead to excessive Ca^2+^ uptake by mitochondria, attenuating their membrane potential, the main driver of ATP production^2^. Hence, energetic failure could have contributed to the inhibition of ATP-driven transport following some experimental manipulations. In light of these findings it is interesting to note that it took ca. 2 minutes to reach the highest level of inhibition following KCl-induced depolarization, but only ca. 30s following the EFS-induced activity patterns. Together, these findings indicate calcium’s role as a regulatory cue is more complex than previously thought, potentially serving a different purpose during realistic firing, than when homeostasis is threatened.

In an effort to study calcium dynamics in the axon, we created a robust method to quantify absolute calcium concentrations with high spatial precision using the ratiometric Ca^2+^-indicator Fura-2. This method was then used to describe the calcium responses in each neuronal compartment following the EFS-induced activity patterns employed during our transport recordings (fig. 4). Axonal calcium levels were observed to scale with firing frequency in the range of 0.1-1µM (fig. 3f). Ratiometric imaging, moreover, did not reveal notable differences in the calcium response amplitude along the length of the axon (i.e. low variability in response amplitude). Hence, the spatial pattern identified in this work, is unlikely to arise due to differences in calcium response amplitude between axonal segments. One potential explanation might be a correlative distribution of syntaphilin^22^, an axon-specific mitochondrial docking protein capable of halting transport in Ca^2+^-dependent manner (i.e. low expression in non-connecting segments).

The main focus of this study was on the axonal compartment. However, ratiometric imaging also provided insights into the calcium dynamics in other neuronal compartments. At rest, a baseline of ca. 50nM was maintained in all compartments, in accordance with previous reports^28–30^. Except in the cytosol, were a higher and more variable baseline was measured of ca. 50-100nM. In ‘baseline maps’ (fig. 8c), produced by an analogue pipeline based on identical calculations performed per pixel instead of per ROI-average signal, these higher baseline values appear to stem from a ‘halo’-like silhouette around the nucleus, reminiscent of the ER^31^. Corrections for background and autofluorescence did not influence this pattern. Since such sharp Ca^2+^ gradients within the cytosolic compartment (or between the cytosolic and dendritic compartments) are implausible, we suspect these halos might arise from dye-leakage into the endoplasmic reticulum. Although this presumed artifact did not interfere with our study, it might present a general issue for quantifications based on cytosolic ROIs using Fura-2. ‘Peak amplitude maps’, generated by the pixel-based pipeline, also revealed a high variability in the calcium responses between dendritic branch segments (fig. 8a). Moreover, this complex pattern was reproduced during repeated stimulations (fig. 8d-i). Local response amplitude, at any given dendritic location, was not correlated to baseline amplitude, nor was it determined by its distance from the soma (fig. 8j-l). These patterns explain the higher variability in dendritic calcium responses observed via the ROI-based approach. Furthermore, the dendritic calcium levels evoked by the activity patterns tested in this study were likely sufficient to trigger transport inhibition, based on previous estimates of its IC50 value^20^. Since branches with high response amplitudes are more likely to halt, and thus accumulate, mitochondria during neuronal activity, these dendritic response patterns might be an important cue to regulate the mitochondrial distribution in the dendritic compartment.

**Figure 8.**
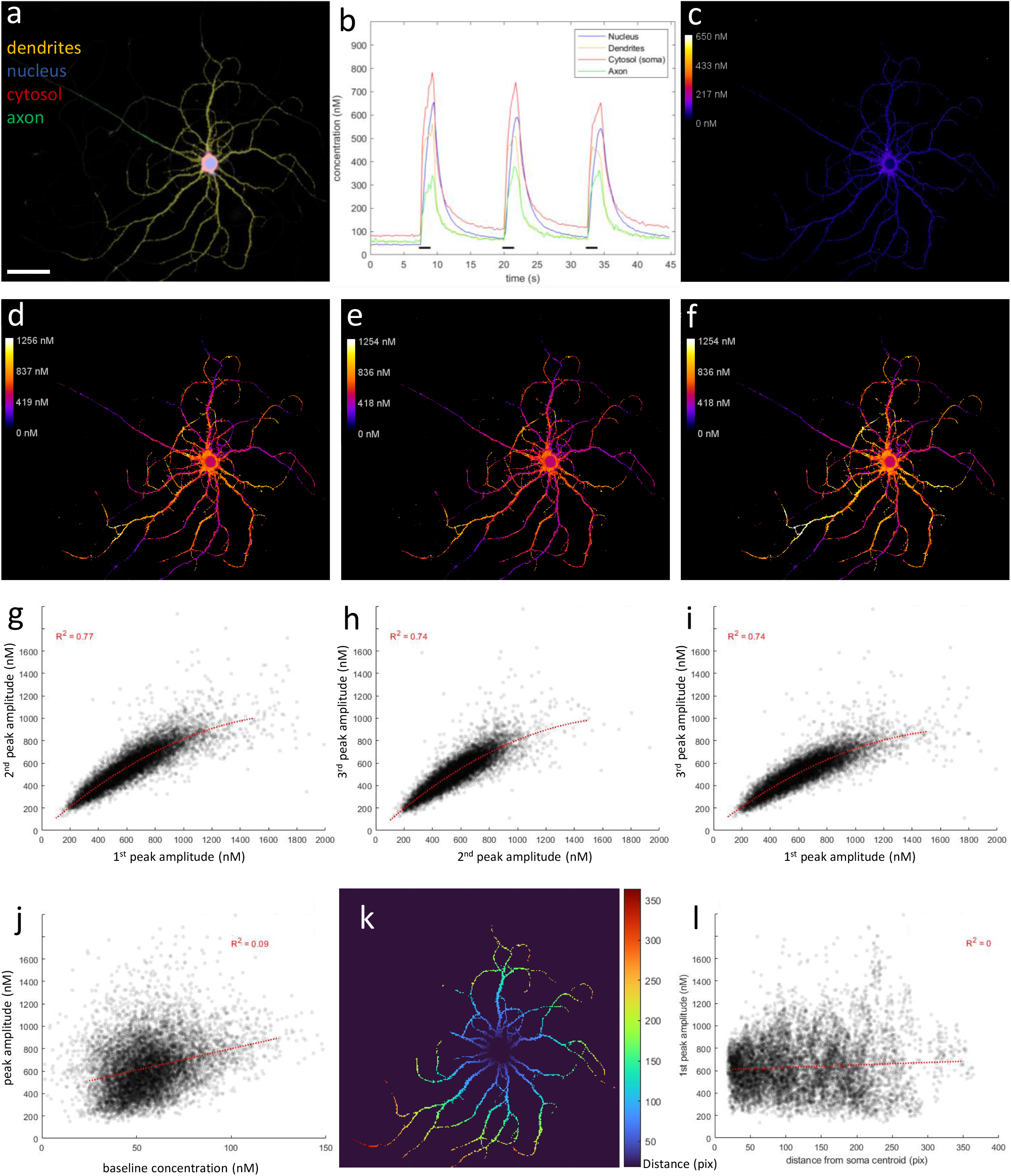
Ratiometric Ca^2+^ imaging reveals complex spatial patterns within dendritic compartment. (a) User-defined compartment ROIs (scalebar = 50µm). (b) Compartment-specific Ca^2+^ responses to repeated 20Hz 2s stimulation (black bars, 3x). (c) Baseline map (average of first 30 frames or 7.5s). A baseline of ca. 50nm is observed in all compartments, except the cytosol, which measured ca. 100nM. (d-f) Peak amplitude maps for each repetition of the stimulation (average of 8 frames during stimulation). Upon stimulation, Ca^2+^ response amplitude is highly variable throughout the dendritic tree. However, this complex spatial pattern is reproduced upon repeated stimulation. (g-i) Scatterplots demonstrating the correlation between response amplitudes during repeated stimulations. (j) Local response amplitudes are not correlated to baseline levels. (k) Distance map, indicating the quasi-Euclidean distance (in pixels) from each ROI pixel to the center of the soma, as the shortest distance traversed within the constraints of the dendritic mask. (l) Local response amplitudes are not correlated to the distance from the soma. Idem for baseline map (data not shown).

## METHODS

### Primary hippocampal cultures

1-2 day old C57BL/6J mouse pups were quickly decapitated before dissection. Hippocampi were dissected in Sylgard dishes containing cold sterile Hank’s buffered salt solution (HBSS in mM: 5.33 KCl, 0.44 KH2PO4, 137.93 NaCl, 0.34Na2HPO40.7H2O, 5.56 D-glucose and 10 HEPES). The tissue was incubated in 0.25% trypsin-ethylenediaminetetraacetic acid (EDTA) (Gibco) supplemented with 200 U/ml DNase (Roche) for 10 min at 37 °C. After three consecutive wash steps with washing buffer (neurobasal medium (Invitrogen) supplemented with 1.59 mg/ml BSA, 0.5% penicillin/streptomycin, 5 mg/ml glucose, 5.5 mg/ml sodium pyruvate, and 200 U/ml DNAaseI) the tissue was mechanically dissociated by trituration. After centrifugation, cells were resuspended and plated on 18 mm diameter coverslips, coated with poly-D-Lysine, suspended over a glial support layer using paraffin dots. The glial support layer was derived using an identical protocol on midbrain and cortical tissue harvest from the same pup. Primary hippocampal neuron-glia co-cultures were grown in a 37 °C, 5% CO2 incubator in Neurobasal-A media (Thermo Fisher Scientific) supplemented with 0.5% penicillin/streptomycin (Lonza), 2% B27 (Gibco), 0.02 mg/ml insulin (Sigma), 50 ng/ml nerve growth factor (Alomone Labs), and 0.5 mM Glutamax (Thermo Fisher Scientific). Half of the medium volume was replaced every 3 days. Imaging was performed after 7-10 days *in vitro*. All procedures were approved by the Animal Ethics Committee of the University of Leuven (Belgium). Ethical Committee number = P110/2020.

### Immunocytochemistry

Coverslips were fixed in 4% PFA for 20min, permeabilized and blocked in 0.5% triton-x solution with host (donkey) serum, labeled with primary antibodies at 4°C overnight (see table below), and finally incubated with secondary antibodies (1:1000) at room temperature for 2h.

| Primary Antibody | Dilution | Purpose | Reference number |
| --- | --- | --- | --- |
| ch-MAP2 | 1:2500-5000 | To define dendritic compartment. | ab5392 (Abcam) |
| g-TAU C17 | 1:500 | To distinguish axonal compartment in combination with MAP2. | Sc-1995 (Santa Cruz) |
| m-ANK G | 1:250 | To define axon initial segment. | Clone N106/20 (Antibodies Inc) |
| rb-TUBB3 | 1:1000 | To distinguish axonal compartment in combination with MAP2. | ab52623 (Abcam) |
| r-alphaTub | 1:500 | To distinguish axonal compartment in combination with MAP2. | MA1-80189 (Thermofisher) |
| r-Syn-1 | 1:500 | Pre-synaptic marker. | ab64581 (Abcam) |

### Axonal mitochondrial transport imaging

Time-lapse imaging of axonal mitochondrial transport and calcium dynamics was performed on a Zeiss LSM 780 confocal microscope, at 37°C in HEPES buffered solution containing in mM; 140 NaCl, 5 KCl, 10 HEPES, 2 CaCl_2_, 2 MgCl_2_ and 10 D-glucose, adjusted to pH = 7.4 using NaOH. Coverslips were loaded with 25nM mitoTracker Red + 25-50nM Fluo-4 for 10 min, washed twice for 5min with HEPES buffer. For fast time-lapse imaging of action potential firing, neurons were loaded with 625nM BeRST1^24^ (gift from Dr. Evan Miller) and 50nM Fluo4. To record mitochondrial transport in ‘connecting’ axonal segments, 10min recordings (0.2Hz) in sparse mixed mito-dendra2:wildtype (50:50) cultures, with manually triggered stimulation after 1min. The connection, as well as axon identity were confirmed using immunolabeling after each recording.

### Stimulation of neurons

At a set time during the recording, electrical field stimulation was applied via a miniature, custom-built bipolar electrode carefully positioned over the field of view, connected to a high current stimulus isolator (WPI, A385) and Master 8 (A.P.I), to generate the desired current amplitude (7.5mA) and stimulus paradigm, respectively. Three stimulation types were used: 20Hz 2sec (40APs), 100Hz 2sec (200APs), 2Hz 2min (240APs).

### Ratiometric calcium imaging

Fura-2 imaging (340/380nm) of neurons was performed on a Nikon inverted widefield microscope equipped with a Polychrome V lamp using TillVision software. Sparse hippocampal cultures were loaded with 0.5µM Fura-2 for 20min at 37°C in plate media, then washed 3x with 37°C HEPES. After a 5min rest, 3 FOVs were measured per coverslip (stimulation types in random order) within a 20min window under continuous perfusion of HEPES buffer at 37°C. After each experiment, MAP2 & TUBB3 labeling was used to distinguish dendritic and axonal compartments. To analyze these recordings, a custom pipeline was created, as described in fig. 3. For the intracellular measurement of maximal (R_max_) and minimal (R_min_) ratios, HEPES buffers containing 5µM ionomycin + 2mM Ca^2+^, and 5µM ionomycin + 0mM Ca^2+^ + 2mM EGTA were used respectively (K_D_ Fura-2 = 228nM).

### Transport analysis

A custom Igor-toolbox was used to generate kymographs and trace mitochondrial trajectories. In house MATLAB scripts were used to analyze transport parameters.

## AUTHOR CONTRIBUTIONS

Experimental work: T.V., Analysis: T.V., P.V.B., Writing & Editing: T.V., P.V.B.

## ACKNOWLEDGEMENTS

The authors’ work is supported by the Research Foundation Flanders (FWO) grant 1111220N. We thank Dr. Evan Miller for providing us with a sample of the BeRST1 voltage sensor.

## DATA AVAILABILITY

The raw recordings that support the findings of this study are available from the corresponding author upon reasonable request. The source data underlying all main and supplementary figures are provided as a Source Data File.

## CODE AVAILABILITY

Custom-written Igor & MATLAB code is available from the corresponding author upon reasonable request.

## REFERENCES

1. Mink, J. W., Blumenschine, R. J. & Adams, D. B. Ratio of central nervous system to body metabolism in vertebrates: its constancy and functional basis. Am. J. Physiol. Integr. Comp. Physiol. 241, R203–R212 (1981).

2. Kann, O. & Kovács, R. Mitochondria and neuronal activity. Am. J. Physiol. Physiol. 292, C641–C657 (2007).

3. Zhu, X.-H. et al. Quantitative imaging of energy expenditure in human brain. Neuroimage 60, 2107–2117 (2012).

4. Attwell, D. & Laughlin, S. B. An Energy Budget for Signaling in the Grey Matter of the Brain. J. Cereb. Blood Flow Metab. 21, 1133–1145 (2001).

5. Vaccaro, V., Devine, M. J., Higgs, N. F. & Kittler, J. T. Miro1-dependent mitochondrial positioning drives the rescaling of presynaptic Ca 2+ signals during homeostatic plasticity. EMBO Rep. 18, 231–240 (2017).

6. Chen, H. & Chan, D. C. Critical dependence of neurons on mitochondrial dynamics. Curr. Opin. Cell Biol. 18, 453–459 (2006).

7. d’Ydewalle, C. et al. HDAC6 inhibitors reverse axonal loss in a mouse model of mutant HSPB1–induced Charcot-Marie-Tooth disease. Nat. Med. 17, 968–974 (2011).

8. Alexander, C. et al. OPA1, encoding a dynamin-related GTPase, is mutated in autosomal dominant optic atrophy linked to chromosome 3q28. Nat. Genet. 26, 211–215 (2000).

9. Civera-Tregón, A. et al. Mitochondria and calcium defects correlate with axonal dysfunction in GDAP1-related Charcot-Marie-Tooth mouse model. Neurobiol. Dis. 152, 105300 (2021).

10. Sheng, Z.-H. & Cai, Q. Mitochondrial transport in neurons: impact on synaptic homeostasis and neurodegeneration. Nat. Rev. Neurosci. 13, 77–93 (2012).

11. Burté, F., Carelli, V., Chinnery, P. F. & Yu-Wai-Man, P. Disturbed mitochondrial dynamics and neurodegenerative disorders. Nat. Rev. Neurol. 11, 11–24 (2015).

12. Kevenaar, J. T. & Hoogenraad, C. C. The axonal cytoskeleton: from organization to function. Front. Mol. Neurosci. 8, 44 (2015).

13. Van Steenbergen, V. et al. Nano-positioning and tubulin conformation contribute to axonal transport regulation of mitochondria along microtubules. Proc. Natl. Acad. Sci. U. S. A. 119, e2203499119 (2022).

14. Sheng, Z.-H. The Interplay of Axonal Energy Homeostasis and Mitochondrial Trafficking and Anchoring. Trends Cell Biol. 27, 403–416 (2017).

15. Vanden Berghe, P., Hennig, G. W. & Smith, T. K. Characteristics of intermittent mitochondrial transport in guinea pig enteric nerve fibers. Am. J. Physiol. Liver Physiol. 286, G671–G682 (2004).

16. Rintoul, G. L., Filiano, A. J., Brocard, J. B., Kress, G. J. & Reynolds, I. J. Glutamate decreases mitochondrial size and movement in primary forebrain neurons. J. Neurosci. 23, 7881–8 (2003).

17. Yi, M., Weaver, D. & Hajnóczky, G. Control of mitochondrial motility and distribution by the calcium signal: a homeostatic circuit. J. Cell Biol. 167, 661–72 (2004).

18. Saotome, M. et al. Bidirectional Ca2+-dependent control of mitochondrial dynamics by the Miro GTPase. Proc. Natl. Acad. Sci. U. S. A. 105, 20728–33 (2008).

19. Wang, X. & Schwarz, T. L. The Mechanism of Ca2+-Dependent Regulation of Kinesin-Mediated Mitochondrial Motility. Cell 136, 163–174 (2009).

20. MacAskill, A. F. et al. Miro1 Is a Calcium Sensor for Glutamate Receptor-Dependent Localization of Mitochondria at Synapses. Neuron 61, 541–555 (2009).

21. Kang, J.-S. et al. Docking of Axonal Mitochondria by Syntaphilin Controls Their Mobility and Affects Short-Term Facilitation. Cell 132, 137–148 (2008).

22. Chen, Y. & Sheng, Z.-H. Kinesin-1–syntaphilin coupling mediates activity-dependent regulation of axonal mitochondrial transport. J Cell Biol 202, 351–364 (2013).

23. Chen, T.-W. et al. Ultrasensitive fluorescent proteins for imaging neuronal activity. Nature 499, 295–300 (2013).

24. Huang, Y.-L., Walker, A. S. & Miller, E. W. A Photostable Silicon Rhodamine Platform for Optical Voltage Sensing. J. Am. Chem. Soc. 137, 10767–10776 (2015).

25. Obashi, K. & Okabe, S. Regulation of mitochondrial dynamics and distribution by synapse position and neuronal activity in the axon. Eur. J. Neurosci. 38, 2350–2363 (2013).

26. Li, Z., Okamoto, K.-I., Hayashi, Y. & Sheng, M. The Importance of Dendritic Mitochondria in the Morphogenesis and Plasticity of Spines and Synapses. Cell 119, 873–887 (2004).

27. Chang, D. T. W., Honick, A. S. & Reynolds, I. J. Mitochondrial trafficking to synapses in cultured primary cortical neurons. J. Neurosci. 26, 7035–45 (2006).

28. Schiller, J. & Sakmann, B. Spatial profile of dendritic calcium transients evoked by action potentials in rat neocortical pyramidal neurones. J. Physiol. 583–600 (1995).

29. Maravall, M., Mainen, Z. F., Sabatini, B. L. & Svoboda, K. Estimating Intracellular Calcium Concentrations and Buffering without Wavelength Ratioing. Biophys. J. 78, 2655–2667 (2000).

30. Ali, F. & Kwan, A. C. Interpreting in vivo calcium signals from neuronal cell bodies, axons, and dendrites: a review. 10.1117/1.NPh.7.1.011402 7, 011402 (2019).

31. de Juan-Sanz, J. et al. Axonal Endoplasmic Reticulum Ca2+ Content Controls Release Probability in CNS Nerve Terminals. Neuron 93, 867–881.e6 (2017).

